# ICOS enhances follicular T helper responses and deteriorates the pathogenic process of liver in mice infected with *Schistosoma japonicum*

**DOI:** 10.1101/2020.02.27.968248

**Authors:** Bo Wang, Song Liang, Yan-Yan Wang, Yu Wang, Chao-Ming Xia

## Abstract

**Background:** Humoral immune responses play an important role in mediating liver granulomatous inflammation and fibrosis in schistosomiasis. Follicular helper T (Tfh) cells have a central role in mediating humoral immune responses. Generation of Tfh cells depends on inducible T cell costimulator (ICOS) signaling, but the underlying molecular mechanisms are incompletely understood in pathogenesis of schistosomiasis.

**Methodology/Principal Findings:** We used a strain of ICOS-transgenic (Tg) mice to test the degrees of liver granulomatous inflammation and fibrosis, the frequency of splenic Tfh cells and soluble egg antigen-specific cytokine responses longitudinally in mice following *Schistosoma japonicum* (*S. japonicum)* infection. In comparison with that in wide-type (WT) mice, significantly severer liver granulomatous inflammation and fibrosis and higher mortality were observed in ICOS-Tg mice. Significantly higher frequency of splenic Tfh cells was accompanied by significantly higher levels of Bcl-6 and CXCR5 expression in the livers of ICOS-Tg mice. Furthermore, significantly higher levels of SEA-specific IL-4, IL-6, IL-10, IL-13, IL-17A, IL-21 and TGF-β1 responses, but lower levels of IFN-γ responses were detected in ICOS-Tg mice, which were abrogated by treatment with ICOS blockers in vitro. In addition, significantly higher levels of serum anti-SEA IgG were detected in ICOS-Tg mice.

**Conclusions/Significance:** The ICOS-related signaling may promote the pathogenesis of murine schistosomiasis by polarizing Tfh cells, which may be immune check points for the prevention and intervention of schistosomiasis.

**Author summary:** Granulomatous inflammation and fibrosis in the liver are the major pathogenic characteristics of schistosomiasis. ICOS is crucial for the development of Tfh cells, which are the key modulators of B cell activation and humoral immunity. However, the underlying molecular mechanisms are incompletely understood in pathogenesis of schistosomiasis. Here, our results showed that the ICOS over-expression would significantly induce severer liver inflammation and fibrosis, higher frequency of splenic Tfh, higher levels of anti-SEA IgG as well as imbalanced SEA-specific cytokine responses in ICOS-Tg mice. The findings suggested that ICOS signaling may promote the pathogenesis of murine schistosoma-related liver inflammation and fibrosis by polarizing Tfh cells. Potentially, ICOS signaling and Tfh cells may be immune check points for the prevention and intervention of schistosomiasis.

## Introduction

*S. japonicum* infection remains a problem for human health in developing countries. During the pathogenesis of schistosomiasis, *S. japonicum* infection-related egg deposition can recruit inflammatory infiltrates and cause granulomatous inflammation and fibrosis in the liver [1,2]. More importantly, continual pathogenic process of inflammation and fibrosis in the liver usually impairs its function, even leading to cirrhosis and live failure. Although previous studies have shown that antigen-specific T cells, such as Th2 and Th17, participate in the pathogenic process and Th1 and regulatory T cells (Tregs) antagonize *Schistosomiasis*-related inflammation [3-5]. The molecular mechanisms of immunoregulation of *S. japonicum* infection-related inflammation and fibrosis in the liver have been incompletely clarified. Hence, illustration of the pathogenic process of *S. japonicum* infection-related inflammation and fibrosis will be of great significance in the develop targets for design of new therapies for patients with schistosomiasis.

Tfh cells are activated CD4^+^ T cells commonly resident in the B cell follicles of second lymph tissues and constitutively express homing receptor CXCR5 [6,7]. Tfh cells can also express programmed death 1 (PD-1), ICOS and CD40L and secrete IL-21 as well as other cytokines, which are crucial for their autocrine regulation of Tfh development and functions [8-10]. Tfh differentiation is mainly regulated by the transcription factor, Bcl-6 [11].Tfh cells can promote antigen-specific B cell activation, germinal center formation and humoral responses [12,13]. In addition, Tfh cells are involved in the pathogenic process of some autoimmune diseases [14-16] and participate in defensing against parasite infection [17,18]. Indeed, increased frequency of Tfh cells is detected in rodent with *S. japonicum* infection and Tfh cells activated by ICOS^+^ macrophages infiltrate into the liver and enhance the liver injury in mice infected with *S. japonicum* [19].

ICOS is crucial for T and B cell activation and important for T and B cell interaction to promote humoral responses [20]. ICOS is highly expressed by human tonsillar CXCR5^+^ T cells within the light zone of germinal centers and efficiently support immunoglobulin production [21,22]. In addition, ICOS-deficient mice show poor humoral responses and ICOS deficiency in humans results in significantly reduced numbers of Tfh cells, indicating a critical role of ICOS in the differentiation of CXCR5^+^ CD4^+^ T cells [23,24]. However, it is incompletely understood that the underlying molecular mechanisms of Tfh development and function in pathogenesis of schistosomiasis. Here we characterized to establish the ICOS-Tg mice as a model of schistosomiasis to assess the role of the ICOSL/ICOS interaction in mediating humoral immune responses by polarizing Tfh cells, particularly, which were abrogated by treatment with ICOS blockers in vitro. ICOS signaling and Tfh cells may be immune check points for the prevention and intervention of schistosomiasis.

## Methods

### Ethics statement

The experimental protocols were established, according to the Regulations for the Administration of Affairs Concerning Experimental Animals (1988.11.1) of the State Science and Technology Commission of the People’s Republic China and were approved by the Institutional Animal Care and Use Committee (IACUC) of Soochow University (Permit Number: 2007-13). All efforts were made to minimize suffering of animals.

### Mice, parasites, and infection

Specific pathogen-free (SPF) female FVB mice (6-8 weeks old) were purchased from the Center of Comparative Medicine of Yangzhou University (Yangzhou, China). Animals were housed and bred in a SPF facility of our campus. To generate ICOS-Tg mice, human ICOS cDNA was amplified by RT-PCR from activated human T cells and cloned into a vector pEGFP-N2 plasmid, and a Tg cassette that drives transgene expression by the CMV promoter/enhancer. The *Ase*I*-Stu*I transgene fragment at 0.642 kbwas microinjected into fertilized mouse eggs prepared from FVB mice.

The generated ICOS-Tg founders were backcrossed to FVB mice over 10 generations to stabilize the Tg strain (Yangzhou, China). Snails (*Oncomelaniahupensis*) harboring *S. japonicum* cercariae (Chinese mainland strain) were purchased from Jiangsu Institute for Schistosomiasis Control (Wuxi, China).

Individual FVB wide-type and ICOS-Tg mice at 6-8 weeks of age were infected with 14 (±1) cercaria of *S. japonicum* through the abdominal skin. At 4, 7, 12, 16, and 20 weeks post-infection, 8 mice at each time point from the infected and control groups were randomly chosen and sacrificed for subsequent *ex vivo* experiments. The remaining 20 mice per group were monitored for their survival until they met certain clinical criteria for sacrifice. The criteria included severe diarrhea, and difficult to eat and breath.

### Flow cytometry

The frequency of Tfh and other subsets of T cells was determined by flow cytometry analysis. Briefly, splenic mononuclear cells were prepared from individual mice at each time point post infection and the cells at 1×10^6^/tube were stained in duplicate with FITC-anti-CD4, PE-anti-CXCR5, PE-Cy5-anti-ICOS, and APC-anti-CD40L or isotype controls (eBioscience, San Diego, USA). The percentages of CXCR5^+^CD4^+^ Tfh cells, ICOS^+^CXCR5^+^CD4^+^ Tfh cells and CD40L^+^CXCR5^+^ CD4^+^ Tfh cells were analyzed on a FACS Calibur Flow Cytometer (BD Biosciences).

In addition, splenic mononuclear cells at 1×10^6^ cells/well were stimulated with 25 ng/ml of phorbolmyristate acetate (PMA) and 1 µg/ml of ionomycin (Sigma-Aldrich) in complete RPMT 1640 medium in the presence of 3 µg/ml of BFA for 5 hours at 37 °C in 5% CO_2_. Subsequently, the cells were surface-stained with FITC-anti-CD4 and PE-anti-CXCR5 or FITC-anti-CD4 alone, fixed, permeabilized with Fixation/Permeabilization buffer, followed by intracellularly staining with APC-anti-IL-21, PE-anti-BCL-6 or isotype controls for analyzing the frequency of IL-21^+^CXCR5^+^CD4^+^ or BCL-6^+^CD4^+^ Tfh cells, respectively.

### Culture and stimulation of splenic mononuclear cells *in vitro*

Splenic mononuclear cells were prepared from individual mice in each group at the indicated time points post infection. Splenic mononuclear cells at 1×10^6^ cells/well were stimulated in triplicate with 25 µg/ml of soluble egg antigens (SEA) of *S. japonicum* from Jiangsu Institute for Schistosomiasis Control (Wuxi, China), 25 ng/ml of PMA and 1 µg/ml of ionomycin in 10% *fetal bovine serum* (FBS) RPMI 1640 in 24-well plates at 37 °C for 72 hours in the presence or absence of 0.5 µg/ml of anti-ICOSL or 0.375 µg/ml of antagonist anti-ICOS (eBioscience). The culture supernatants were collected and the contents of cytokines (IL-2, IL-4, IL-6, IL-10, IL-13, IL-17A, IL-21, IFN-γ, TNF-α) were determined by Cytometric Bead Array (CBA) using specific kit (BD Biosicences, San Diego, USA), according to the manufacturers’ instruction. The levels of TGF-β1in the cultured supernatants were measured by ELISA using the ELISA Ready-SET-Go kit (eBiosicence, San Diego, CA), according to the manufacturers’ protocol. The limitation of detection for IL-2, IL-4, IL-6, IL-10, IL-13, IL-17A, IL-21, IFN-γ, TNF-α and TGF-β1 is (2.8pg/ml) respectively.

### Enzyme-linked immunosorbent assay (ELISA)

The concentrations of serum SEA-specific IgG and HA in individual mice were evaluated by ELISA. Briefly, individual wells of 96-well ELISA plates (ICN Biomedicals, Costa Mesa, USA) were coated with 100 µl of SEA (20 µg/ml) in 0.05 M carbonate buffer (pH 9.6) overnight at 4°C and after being washed and blocked with 10% FCS in 0.01M PBST, the wells were added in triplicate with 1:100 diluted serum samples and cultured at 37 °C for 1 hour. The bound antibodies were detected with horseradish peroxidase (HRP)-conjugated goat anti-mouse IgG (1:4000) for 1 hour at 37 °C and developed with tetramethyubenzidine (TMB) substrate (BD PharMigen), followed by measuring the absorbance of individual wells at 450 nm using an ELISA reader (Bio-rad mod.550). Serum samples from unmanipualted mice were used for negative controls.

### Histopathological study

The liver tissue samples from individual mice were dissected out at the indicated time points post infection, fixed with paraformaldehyde and paraffin-embedded. The tissue sections at 5 μm were stained with hematoxylin-eosin and the egg-related granuloma volumes of individual liver samples were evaluated using a formula of V=πAB^2^/6, where A and B represented two diameters cross opposite axes [5].

Furthermore, some liver tissue sections were stained with Masson trichrome (MT). The degrees of liver fibrosis in at least 10 low-power fields (magnification x 100) of each sample were evaluated by pathologists in a blinded manner. In addition, the percentages of collagen+ regions were calculated using an image analysis system (Image-Pro Plus 6.0) to measure a relative objective index. The fibrotic areas were calculated and expressed as the relative ratios of the collagen-containing area to the whole area. At least 10 high-power fields (magnification x 200) were measured from each liver sample.

### Immunohistochemistry

The contents of CXCR5 and Bcl-6 expression in the livers of individual mice were evaluated by immunohistochemistry. Briefly, liver tissue sections (5 µm) were rehydrated, and treated with 3% H_2_O_2_, followed by blocking with 10% normal goat serum (Boster, Wuhan, China). Subsequently, the sections were incubated with rabbit anti-CXCR5 (1:500, Millipore, Germany) or rabbit anti-Bcl-6 (1:100, Santa Cruz, USA) overnight at 4 °C and bound antibodies were detected using HRP-conjugated goat anti-rabbit IgG (ChemMate Envision/HRP, rabbit/mouse detection kit, Gene-tech, Shanghai, China) at 37 °C for 1 hour. After being washed, the immune staining was visualized using 3, 3’-Diaminobenzidine (DAB) and counterstained with Mayer’s hematoxylin. Rabbit sera at 1:10 dilutions served as negative controls. The intensity and percentages of positive staining cells in the liver granulomas were evaluated for five high-power fields selected randomly using the Leica QWin Plus software, version 3.5.1 (Leica Microsystems, Switzerland). The percentage of positively staining cells in 15 granulomas was assessed and the mean of % positively staining cells in 15 granulomas +/- SD in each group was calculated.

### Statistical analysis

Statistical analysis was performed using the SPSS version 10.1 (Statistical Package for Social Science, Chicago, IL) software. The difference of the data among the groups was compared by two-way ANOVA followed by Tukey’s multiple comparison test. The relationship between measures was analyzed by Spearman’s rank correlation. The survival of each group of mice was estimated by Kaplan-Meier survival analysis and the difference between two groups was analyzed by the log-rank test. A P value of P<0.05 were considered statistically significant.

## Results

### ICOS over-expression deteriorates the liver granulomatous inflammation and fibrosis in mice post-infection

*S. japonicum* infection can cause liver granulomatous inflammation and fibrosis. To assess the impact of ICOS over-expression on *S. japonicum* infection-related liver inflammation the liver and spleen gross pathology, weights and liver granulomatous inflammation were examined longitudinally in WT and ICOS-Tg mice following *S. japonicum* infection. We observed dark-yellow livers (Fig 1A-b,c) compared with non-infected liver (Fig 1A-a) and enlarged spleens in the mice infected with *S. japonicum* (Fig 1A-d). The liver and spleen weights in the mice infected with *S. japonicum* rapid increased and significantly greater than that in the non-infected mice (Fig 1B). The liver and spleen weights in ICOS-Tg mice were significantly greater than that in the WT mice at 4 weeks post infection and later time points (Fig 1B; \**P*<0.05 or \*\**P*<0.01). Histological examination indicated that the mean volumes of granulomas in the ICOS-Tg mice were also significantly bigger than that in the WT mice (Fig 1C and D). More importantly, the overall survival periods of ICOS-Tg mice were significantly shorter than that in the WT mice (Fig 1E; *P*<0.01). Collectively, these data indicated that induction of systemic ICOS over-expression deteriorated the liver granulomatous inflammation and fibrosis and accelerated mortality in mice infected with *S. japonicum*.

**Fig 1.**
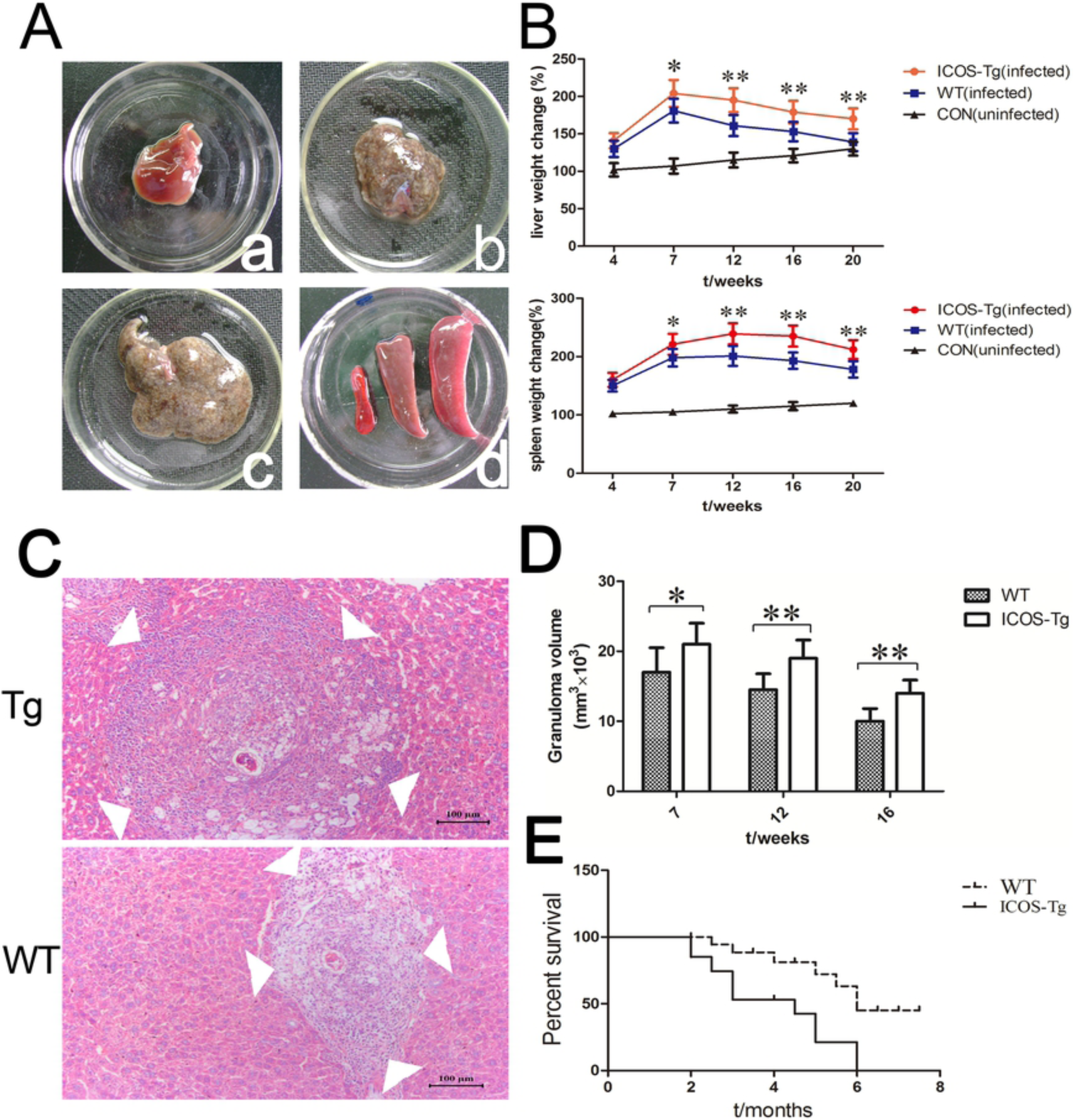
Hepatic granulomatous inflammation and fibrosis in ICOS-Tg mice post-infection. Female FVB ICOS-Tg and WT mice were infected with *S. japonicum*. The liver and spleen gross pathology, weights and liver granulomatous inflammation in WT and ICOS-Tg mice were examined longitudinally at the indicated time points post *S. japonicum* infection. The liver tissue sections were stained with H&E and granulomous inflammatory areas were analyzed in a blinded manner. Data are representative images and expressed as the mean ± SEM of each group (n=8 per group per time point). The survival of remaining mice (n=20 per group) were monitored. (**A**) The gross pathology of the livers from the healthy control (**a**); infected WT mice (**b**) and ICOS-Tg mice (**c**) and spleens (**d**: from left to right: normal spleen; infected WT mice; infected ICOS-Tg mice). (**B**) The changes in the liver and spleen weights. The liver and spleen weights of healthy mice at 4 weeks post infection was designated as 100%. Two-way ANOVA followed by Tukey’s multiple comparison test showed no significant genotype × time interaction effect for the liver/spleen weight changes (*P*=0.455/ *P*=0.16). The main effects showed a significant difference in the changes of the liver/spleen weights among different time points and different genotypes (\*\**P*<0.01). (**C**) H&E analysis of the liver sections of mice at 12 weeks post infection. The images are original magnification, ×100; scale bar, 100um. Arrowheads indicate the margins of hepatic granulomatous inflammation. (**D**) The mean granuloma sizes in ICOS-Tg and WT mice were assessed at 7, 12, or 16 weeks post-infection and quantified from at least 20 granulomas selected randomly from individual mice, as described in the Materials and Methods. \**P*<0.05, \*\**P*<0.01, or \*\*\**P*<0.001 vs. the WT mice. Two-way ANOVA and then Tukey’s multiple comparison tests showed a significant genotype × time interaction effect for the volumes of egg granuloma (\**P*<0.05). The main effects showed a significant difference in the volumes of egg granuloma among different time points and different genotypes (\**P*<0.05 or \*\**P*<0.01). (**E**) The survival of mice following *S. japonicum* infection. (ICOS-Tg vs. WT, *P*=0.0038). The survival analysis data are a representative of three independent experiments with similar results.

Given that chronic *S. japonicum* infection is associated with the liver fibrosis we examined the effects of systemic ICOS over-expression on the liver fibrosis in mice infected with *S. japonicum*. MT staining revealed that the liver fibrosis developed at 7 weeks post infection and the degrees of liver fibrosis increased with time in mice following *S. japonicum* infection (Fig 2A). Quantitative analysis pointed out that the degrees of liver fibrosis in the ICOS-Tg mice were significantly greater than that in the WT mice (Fig 2B; *P*<0.01). Therefore, induction of systemic ICOS over-expression enhanced the fibrotic process in the livers of mice following *S. japonicum* infection.

**Fig 2.**
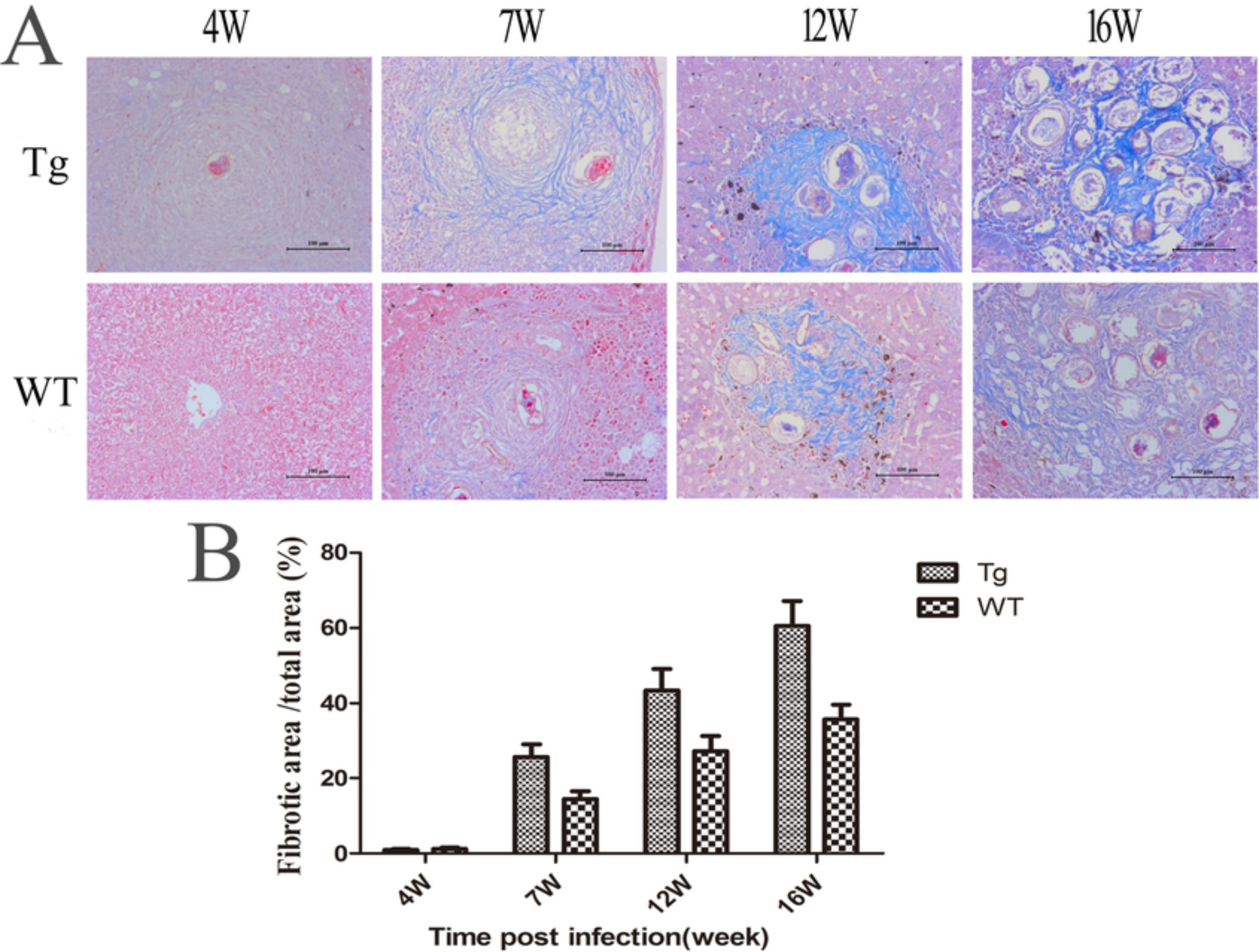
The schistosoma-related liver fibrosis in ICOS-Tg mice. The liver tissue sections from individual mice at the indicated time points post infection were stained with MT (original magnification×200). (**A**) The blue area represents fibrillar collagen. (**B**) Two-way ANOVA and Tukey’s multiple comparison tests showed a significant genotype × time interaction effect for the ratios of fibrotic to total area (%) (\*\**P*=0.0035). The main effects showed a significant difference in the ratios of fibrotic to total area (%) among different time points and different genotypes (\*\**P*<0.01).

### ICOS over-expression increases the frequency of circulating Tfh cells in mice post-infection

CXCR5^+^Bcl-6^+^Tfh cells can secrete IL-21 and are crucial for humoral responses [9]. Recent studies indicate that ICOS positively regulates the Tfh development [8]. To understand the mechanisms underlying the action of ICOS over-expression in this model, we characterized the frequency of circulating Bcl-6^+^CD4^+^, CXCR5^+^CD4^+^ and CXCR5^+^IL-21^+^ T cells in total CD4^+^ T cells in WT and ICOS-Tg mice at different time points post infection by flow cytometry analysis (Fig 3). Quantitative analysis indicated that the percentages of Bcl-6^+^CD4^+^, CXCR5^+^CD4^+^ and CXCR5^+^IL-21^+^ Tfh cells in both strains of mice gradually increased, as compared with that before infection and peaked at 12 weeks post infection, followed by slightly declined. Interestingly, the percentages of Bcl-6^+^CD4^+^, CXCR5^+^CD4^+^ and CXCR5^+^IL-21^+^ Tfh cells in the ICOS-Tg mice were significantly higher than that in the WT mice. Similarly, further analysis revealed that the percentages of circulating ICOS^+^ CXCR5^+^CD4^+^ and CD40L^+^CXCR5^+^CD4^+^ Tfh cells in the ICOS-Tg mice were significantly higher than that in the WT mice (Fig 4). Hence, induction of systemic ICOS over-expression enhanced Tfh responses in mice following *S. japonicum* infection.

**Fig 3.**
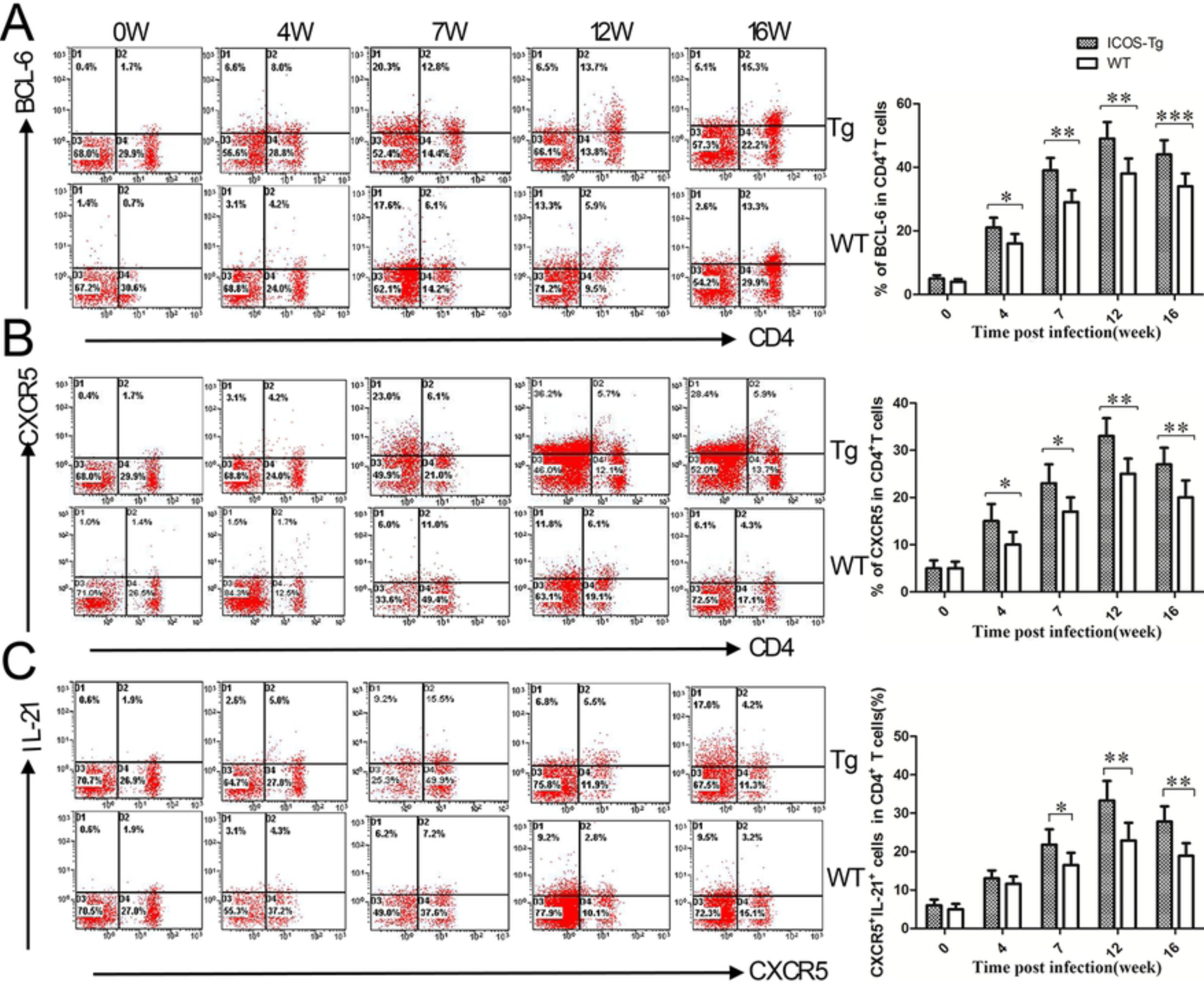
The frequency of splenic Tfh cells. The frequency of splenic Bcl-6^+^CD4^+^ (**A**), CXCR5^+^CD4^+^ (**B**) and IL-21^+^CXCR5^+^ (**C**) Tfh cells in the ICOS-Tg and WT mice was characterized longitudinally post infection by flow cytometry after staining with the indicated antibodies. The cells were first gated on splenic mononuclear cells and at least 10,000 events from each sample were analyzed. Data are representative charts and expressed as the mean ± SEM of the percentages of specific type of Tfh cells in each group (n=5 per group per time point) at the indicated time points post infection from three separate experiments.\**P*<0.05, \*\**P*<0.01, or \*\*\**P*<0.001 vs. the WT mice. Two-way ANOVA and Tukey’s multiple comparison tests showed a significant genotype × time interaction effect for the Bcl-6 (\**P*=0.028), CXCR5 (\**P*=0.033) and IL-21 (\**P*=0.016). The main effects showed a significant difference in the levels of Bcl-6, CXCR5 and IL-21 expression among different genotypes and different times point (\**P*<0.05 or \*\**P*<0.01).

**Fig 4.**
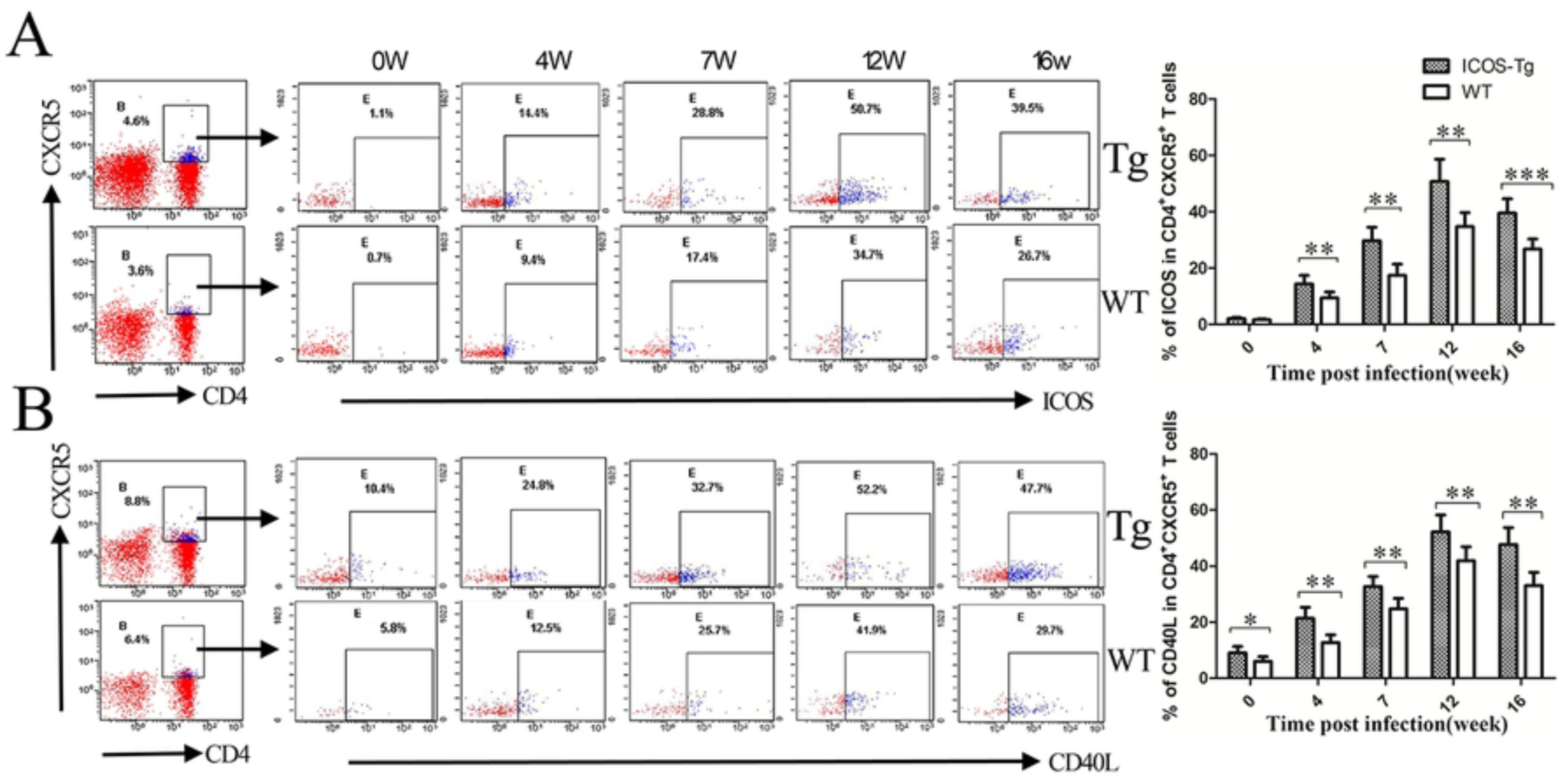
The frequency of splenic ICOS^+^ or CD40L^+^ Tfh cells. The frequency of splenic ICOS^+^CXCR5^+^CD4^+^ Tfh cells and CD40L^+^CXCR5^+^CD4^+^ Tfh cells in the ICOS-Tg and WT mice was characterized by flow cytometry analysis and at least 10,000 events were analyzed. Data are presentative charts and expressed as the mean ± SEM of the percentages of ICOS^+^CXCR5^+^CD4^+^ Tfh cells (**A**) and CD40L^+^CXCR5^+^CD4^+^ Tfh cells (**B**) in each group (n=5 per group per time point) at the indicated time points post infection from three separate experiments. \**P*<0.05, \*\**P*<0.01, or \*\*\**P*<0.001 vs. the WT mice. Two-way ANOVA and Tukey’s multiple comparison tests showed a significant genotype × time interaction effect for the ICOS^+^ Tfh cells (\*\**P*=0.001) and CD40L^+^ Tfh cells (\**P*=0.049). The main effects showed a significant difference in the percentages of ICOS^+^ Tfh cells and CD40L^+^ Tfh cells among different genotypes and different times point (\**P*<0.05 or\*\**P*<0.01).

### ICOS over-expression promotes antigen-specific Tfh, Th2 and Th17 responses in mice post-infection

To further elucidate the role of ICOS-ICOSL signaling in modulating Tfh responses, we isolated splenic mononuclear cells from the ICOS-Tg and WT mice at different time points post *S. japonicum* infection and stimulated them with SEA in the presence or absence of anti-ICOS or anti-ICOSL in vitro, followed by measuring cytokines in the supernatants by CBA. Although the levels of IL-2 and TNF-α increased for a short period post infection in both groups of mice, there was no significant difference in the levels of IL-2 between the ICOS-Tg and WT mice regardless the presence or absence of anti-ICOS or anti-ICOSL (Fig 5A and B). Similarly, the levels of IFN-γ also increased for a short period post infection. The levels of IFN-γ in the supernatants of cultured splenic mononuclear cells from the ICOS-Tg mice were significantly lower than that in the WT mice (Fig 5C). The levels of IFN-γ increased dramatically after treatment with anti-ICOS or anti-ICOSL in the supernatants of cultured splenic mononuclear cells from the ICOS-Tg mice, but not from WT mice in most time points tested. Furthermore, the levels of IL-4, IL-10, IL-13, IL-17A, IL-21 and TGF-β1 gradually increased in both groups of mice, peaked at 12 weeks post infection and slightly declined (Fig 5D-I). The levels of IL-4, IL-10, IL-13, IL-17A, IL-21 and TGF-β1 in the supernatants of cultured splenic mononuclear cells from the ICOS Tg mice were significantly higher than that in the WT mice. Similarly, treatment with anti-ICOS or anti-ICOSL also significantly reduced the levels of IL-4, IL-10, IL-13, IL-17A, IL-21 and TGF-β1 in the supernatants of cultured splenic mononuclear cells from the ICOS Tg mice (Fig 5D-I). These data clearly demonstrated that induction of systemic ICOS over-expression promoted Tfh, Th2 and Th17 responses in mice infection with *S. japonicum.*

**Fig 5.**
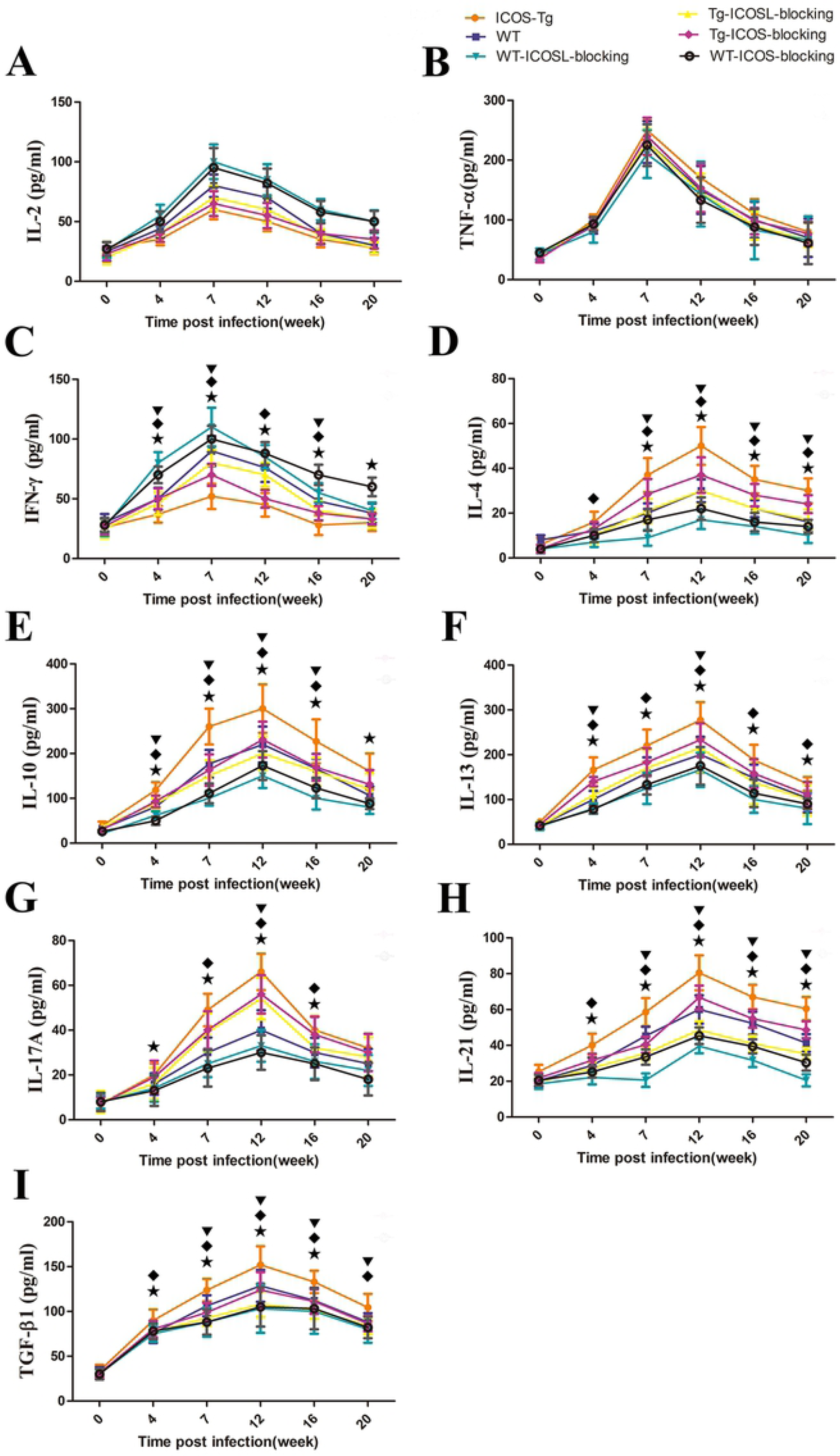
The dynamic changes in the cytokine production by splenocytes from the ICOS-Tg and WT mice. Female ICOS-Tg and WT mice were infected with *S. japonicum*. Splenic mononuclear cells were prepared from each group before and at 4, 7, 12, 16, or 20 weeks post-infection and stimulated in triplicate with SEA, PMA and ionomycin in the presence or absence of anti-ICOS or anti-ICOSL for 72 hrs. The supernatants were harvested and the concentrations of (**A**) IL-2, (**B**) TNF-α, (**C**) IFN-γ, (**D**) IL-4, (**E**) IL-10, (**F**) IL-13, (**G**) IL-17A, (**H**) IL-21 and (**I**) TGF-β1 were determined by CBA. Data are expressed as the mean ± SEM of the concentrations of each cytokine in individual group of mice (n=5 per group, per time point) from three independent experiments. ^¾^*P*<0.05 between ICOS-Tg and the WT mice, ^1^ *P*<0.05 between ICOS-Tg and Tg-ICOSL-blocking mice, ^▾^*P*<0.05 between ICOS-Tg and Tg-ICOS-blocking mice. Two-way ANOVA and Tukey’s multiple comparison tests showed a significant genotype × time interaction effect for the IFN-γ (\**P*=0.046), IL-4 (\*\**P*=0.005), IL-10 (\**P*=0.031), IL-13 (\*\**P*<0.01), IL-17A (\**P*=0.038), IL-21 (\**P*=0.028) and TGF-β1 (\*\**P*=0.004). The main effects showed a significant difference among different genotypes and different time points in the level of IL-2, TNF-α, IFN-γ, IL-4, IL-10, IL-13, IL-17A, IL-21 and TGF-β1 (\**P*<0.05 or \*\**P*<0.01).

Then, we determined the HA titers in ICOS-Tg mice following *S. japonicum* infection and analyzed the potential correlation between the levels of HA and the levels of serum IL-4, IL-13, TGF-β1, and IL-21 (Fig 6A-D). The levels of serum IL-4, IL-13, and TGF-β1 were positively correlated with the HA titers in ICOS-Tg mice following *S. japonicum* infection (Fig 6E; *P*<0.01 or *P*<0.001). Importantly, the levels of serum IL-21 were also positively correlated with the HA titers in ICOS Tg mice following *S. japonicum* infection (*P*<0.05) (Fig 6D and E). These data suggest that ICOSL/ICOS interactions play a key role in Tfh, Th2 and Th17 responses, which are identified as pertinent to fibrosis.

**Fig 6.**
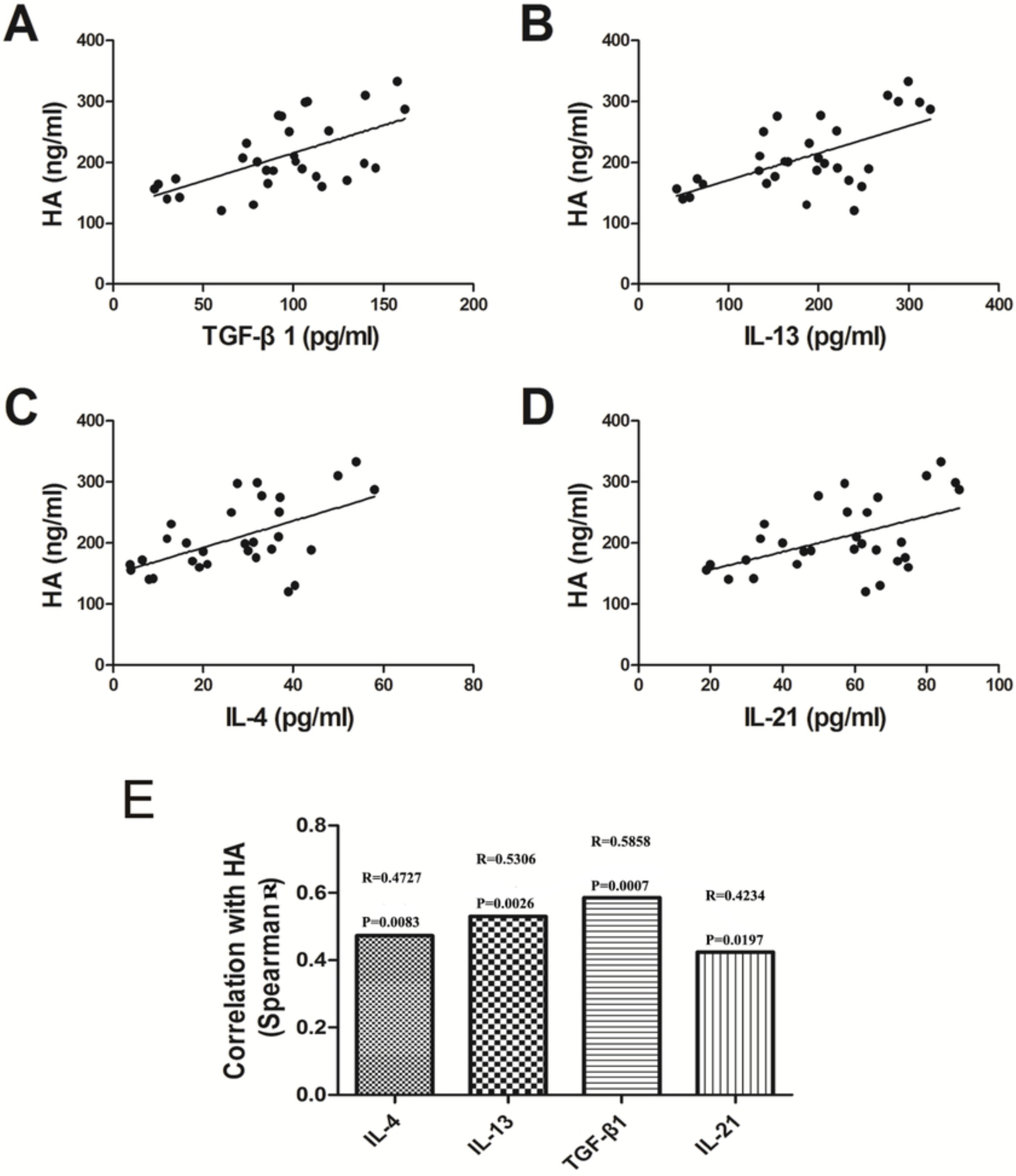
The analysis of correlation of Th2 / Tfh and fibrosis. The linear relationship of IL-4 (**A**: R=0.4727, \*\**P*=0.0083), IL-13 (**B**: R=0.5306, \*\**P*=0.0026), TGF-β1 (**C**: R=0.5858, \*\*\**P*=0.0007), IL-21 (**D**: R=0.4234, \**P*=0.0197) and HA. The results are a representative of three independent experiments with similar results, which are from 6 time points (at 0, 4, 7, 12, 16, 20 weeks) (**E**). The comparison of Spearman R from the analysis of correlation. The Spearman R from the analysis of correlation between IL-4 / IL-13 / TGF-β1 / IL-21 and HA. \**P*<0.05, \*\**P*<0.01, and \*\*\**P*<0.001 between cytokine and HA, Spearman’s rank.

### ICOS over-expression enhances antigen-specific humoral responses and Tfh liver infiltration in mice post-infection

Tfh cells can promote humoral response. Finally, we assessed the impact of systemic ICOS over-expression on SEA-specific humoral responses and on the degrees of liver Tfh infiltration in mice at different time points post *S. japonicum* infection. We found that *S. japonicum* infection induced SEA-specific IgG responses, which gradually increased and peaked at 12 weeks post infection in both groups of mice. The levels of serum SEA-specific IgG in the ICOS-Tg mice were significantly higher than that in the WT mice at chronic stage of *S. japonicum* infection (Fig 7A). Immunohistochemistry analyses indicated that the intensity and extensity of positively anti-BCL-6 and anti-CXCR5 staining in the liver sections of ICOS-Tg mice were obviously stronger than that in the WT mice at 12 weeks post infection (Fig 7B). Quantitative analysis revealed that the ratios of IOD to AOVI for anti-Bcl-6 and anti-CXCR5 staining in the livers of ICOS Tg mice were significantly greater than that in the WT mice at 12 weeks post infection (Fig 7C). Therefore, these data clearly indicated that induction of systemic ICOS over-expression promoted antigen-specific humoral responses and Tfh liver infiltration in mice following *S. japonicum* infection.

**Fig 7.**
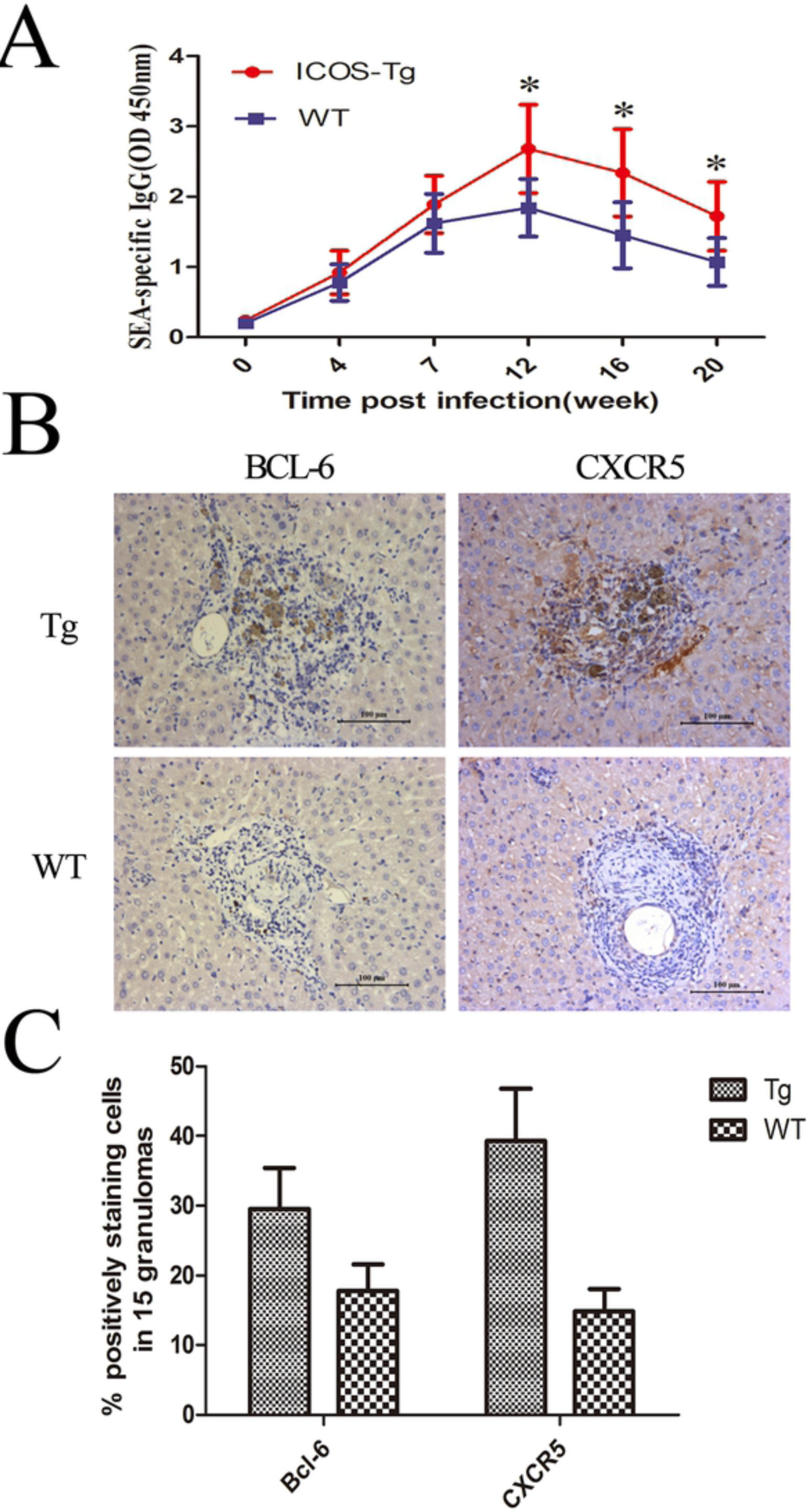
The concentrations of serum SEA-specific IgG and the intensity of Tfh liver infiltration liver in mice. The concentrations of serum anti-SEA IgG in the ICOS-Tg and WT mice at the indicated time points post infection were determined by ELISA (**A**) and the degrees of Tfh infiltrates in the livers of both groups of mice at 12 weeks post infection were determined by immunohistochemistry using anti-Bcl-6 and anti-CXCR5 (**B**). The intensity of anti-Bcl-6 and anti-CXCR5 staining was analyzed by the % positively staining cells in 15 granulomas (**C**). Data are representative images and expressed as the mean ± SEM of the concentrations of serum IgG and the mean of % positively staining cells in 15 granulomas +/- SD of each group of mice (n=5 per group per time point) from three separate experiments. \**P*<0.05, \*\**P*<0.01, or \*\*\**P*<0.001 vs. the WT mice. Two-way ANOVA and Tukey’s multiple comparison tests showed no significant genotype × time interaction effect for the concentrations of serum anti-SEA IgG (*P*=0.08). The main effects showed a significant difference in the changes of the liver/spleen weights among different time points and different genotypes (\*\**P*<0.01).

## Discussion

Tfh cells are crucial for B cell activation, germinal center formation and humoral responses [7]. ICOS, besides CD40L and CD28, is important for T cell activation, particularly for Tfh cells. In this study, we tested the impact of systemic ICOS over-expression on the dynamics of liver granulomatous inflammation and fibrosis in mice following *S. japonicum* infection. We found that in comparison with that in the WT mice, larger liver and spleens organs and greater wet liver and spleen weights were observed, accompanied by significantly severer liver granulomatous inflammation and fibrosis in the ICOS-Tg mice. As a result, the ICOS-Tg mice had significantly shorter survival following *S. japonicum* infection. Clearly, these data demonstrated that systemic ICOS over-expression promoted the liver granulomatous inflammation, fibrosis and mortality in mice following *S. japonicum* infection. These novel findings indicated that systemic ICOS over-expression not only promoted the liver granulomatous inflammation and fibrosis, but also accelerated the mortality in *S. japonicum*-infected mice.

Recent studies have shown that the ICOS signaling can promote Tfh cell differentiation and IL-21 production [9,10]. To understand the mechanisms underlying the action of ICOS over-expression, we characterized the frequency of splenic Tfh cells in the ICOS-Tg and WT mice following *S. japonicum* infection. We found that the percentages of Bcl-6^+^CD4^+^, CXCR5^+^CD4^+^, CXCR5^+^IL-21^+^ and CD40L^+^CXCR5^+^CD4^+^ Tfh cells gradually increased in both groups of mice following *S. japonicum* infection and the percentages of these Tfh cells in the ICOS-Tg mice were significantly higher than that in the WT mice. These data indicated that ICOS enahcned Tfh development during the process of *S. japonicum* infection. It is possible that ICOS may up-regulate the CXCR5 and Bcl-6 expression to promote the differentiation of Tfh cells. More importantly, we observed significantly higher levels of IL-21 secreted by splenic mononuclear cells from ICOS-Tg mice following *S. japonicum* infection. Given that IL-21 is a growth factor and important for the function of Tfh cells, it is possible that the ICOS signaling may also enhance IL-21 expression, providing a positive feedback regulation of Tfh responses. The higher frequency of Tfh cells, the higher levels of IL-21 production may activate and expand memory B cells, thereby enhancing humoral responses. Indeed, we detected significantly higher levels of SEA-specific humoral responses in the ICOS-Tg mice. Apparently, ICOS through promoting Tfh development and in turn enhancing humoral responses, contributes to the pathogenesis of granulomatous inflammation and fibrosis in mice following *S. japonicum* infection. Potentially, ICOS and Tfh may be immune check points for the prevention and intervention of *Schistosoma-*related granulomatous inflammation and fibrosis.

It is well known that Th2 and Th17 responses are involved in the pathogenesis of *Schistosoma-*related granulomatous inflammation and fibrosis. In this study, we found that significantly higher levels of SEA-specific Th2 and Th17 responses in the ICOS-Tg mice, which was abrogated by treatment with a blocker of the ICOS/ICOSL signaling. These data suggest that the ICOS signaling may promote Th2 and Th17 responses during the pathogenic process of *Schistosoma-*related granulomatous inflammation and fibrosis. Our findings were consistent with previous observations that the ICOS expands Th2 immunity and Th2-medaited inflammation by augmenting the migration of inflammatory infiltrates [25,26]. Interestingly, we also detected significantly higher levels of IL-10 and TGF-β1 responses in the ICOS-Tg mice. Given that IL-10 and TGF-β1 are important anti-inflammatory cytokines secreted by regulatory T cells (Tregs), the enhanced anti-inflammatory responses may inhibit the liver granulomatous inflammation. Hence, the ICOS signaling has dual functions by promoting pathogenic Th2 and Th17 responses and enhancing anti-inflammatory IL-10 and TGF-β1 responses during the pathogenesis of *Schistosoma-*related granulomatous inflammation. Unfortunately, TGF-β1 is potent pro-fibrotic factor and through its receptor to activate the Smad signaling to promote fibrosis-related gene expression [27,28]. It is possible that the ICOS signaling through enhancing TGF-β1 production contributes to the pathogenesis of *Schistosoma-*related liver fibrosis. Therefore, inhibition of the ICOS signaling may not only attenuate hepatic granulomatous inflammation, but also mitigate *Schistosoma-*related liver fibrosis.

In summary, our data indicated that systemic ICOS over-expression deteriorated the pathogenic process of *Schistosoma-*related liver granulomatous inflammation and fibrosis and accelerated mortality in mice. Furthermore, systemic ICOS over-expression not only significantly increased the frequency of splenic Tfh cells, but also enhanced SEA-specific Th2, Th17, IgG, IL-21, IL-10 and TGF-β1 responses in mice following *S. japonicum* infection. Hence, the ICOS signaling has sequential roles in regulating the *Schistosoma*-related liver granulomatous inflammation and fibrosis in mice. Importantly, the ICOS signaling may be an immune check point for the prevention and intervention of *Schistosoma-*related liver granulomatous inflammation and fibrosis. Therefore, our findings may not only provide new insights into the mechanisms by which the ICOS regulates the pathogenesis of *Schistosoma-*related liver granulomatous inflammation and fibrosis, but also may aid in the design of new therapies for the intervention of *Schistosoma*-induced fibrosis.

## Notes

**Funding:** This work was supported by grants from the National Natural Science Foundation of China (No: 81171603 and No: 81471977) to C.M.X and the General Program of Inner Mongolia Natural Science Foundation of China (No:2016BS0812), A project funded by the Program of Inner Mongolia People’s Hospital Foundation (No:2019YN23) to B.W. The funders had no role in study design, data collection and analysis, decision to publish, or preparation of the manuscript.

## References

1. Nono JK, Ndlovu H, Aziz NA, Mpotje T, Hlaka L, Brombacher F. Host regulation of liver fibroproliferative pathology during experimental schistosomiasis via interleukin-4 receptor alpha. PLoS Negl Trop Dis. 2017 Aug 21;11(8):e0005861. doi: 10.1371/journal.pntd.0005861. PMID: 28827803; PMCID: PMC5578697.

2. Vicentino ARR, Carneiro VC, Allonso D, Guilherme RF, Benjamim CF, Vicentino ARR, et al. Emerging Role of HMGB1 in the pathogenesis of schistosomiasis liver fibrosis. Front Immunol. 2018 Sep 12;9:1979. doi: 10.3389/fimmu.2018.01979. PMID: 30258438; PMCID: PMC6143665.

3. Kumar R, Mickael C, Kassa B, Sanders L, Koyanagi D, Hernandez-Saavedra D, et al. Th2 CD4+ T Cells are necessary and sufficient for Schistosoma-pulmonary hypertension. J Am Heart Assoc. 2019 Aug 6;8(15):e013111. doi: 10.1161/JAHA.119.013111. Epub 2019 Jul 24. PMID: 31339057; PMCID: PMC6761627.

4. Soloviova K, Fox EC, Dalton JP, Caffrey CR, Davies SJ. A secreted schistosome cathepsin B1 cysteine protease and acute schistosome infection induce a transient T helper 17 response. PLoS Negl Trop Dis. 2019 Jan 17;13(1):e0007070. doi: 10.1371/journal.pntd.0007070. PMID: 30653492; PMCID: PMC6353221.

5. Wang B, Liang S, Wang Y, Zhu XQ, Gong W, Zhang HQ, et al. Th17 down-regulation is involved in reduced progression of schistosomiasis fibrosis in ICOSL KO mice. PLoS Negl Trop Dis. 2015 Jan 15;9(1):e0003434. doi: 10.1371/journal.pntd.0003434. PMID: 25590646; PMCID: PMC4295877.

6. Qin L, Waseem TC, Sahoo A, Bieerkehazhi S, Zhou H, Galkina EV, et al. Insights into the molecular mechanisms of T follicular helper-mediated immunity and pathology. Front Immunol. 2018 Aug 15;9:1884. doi: 10.3389/fimmu.2018.01884. PMID: 30158933; PMCID: PMC6104131.

7. Wu H, Deng Y, Zhao M, Zhang J, Zheng M, Chen G, et al. Molecular control of follicular helper T cell development and differentiation. Front Immunol. 2018 Oct 25;9:2470. doi: 10.3389/fimmu.2018.02470. PMID: 30410493; PMCID: PMC6209674.

8. Uwadiae FI, Pyle CJ, Walker SA, Lloyd CM, Harker JA. Targeting the ICOS/ICOS-L pathway in a mouse model of established allergic asthma disrupts T follicular helper cell responses and ameliorates disease. Allergy. 2019 Apr;74(4):650–662. doi: 10.1111/all.13602. Epub 2018 Nov 12. PMID: 30220084; PMCID: PMC6492018.

9. Wang Y, Lin C, Cao Y, Duan Z, Guan Z, Xu J, et al. Up-regulation of interleukin-21 contributes to liver pathology of schistosomiasis by driving GC immune responses and activating HSCs in mice. Sci Rep. 2017 Nov 30;7(1):16682. doi: 10.1038/s41598-017-16783-7. PMID: 29192177; PMCID: PMC5709429.

10. Shi J, Hou S, Fang Q, Liu X, Liu X, Qi H. PD-1 controls follicular T helper cell positioning and function. Immunity. 2018 Aug 21;49(2):264-274.e4. doi: 10.1016/j.immuni.2018.06.012. Epub 2018 Jul 31. PMID: 30076099; PMCID: PMC6104813.

11. Read KA, Powell MD, Oestreich KJ. T follicular helper cell programming by cytokine-mediated events. Immunology. 2016 Nov;149(3):253–261. doi:10.1111/imm.12648. Epub 2016 Aug 16. PMID: 27442976; PMCID: PMC5046059.

12. Kim ST, Choi JY, Lainez B, Schulz VP, Karas DE, Baum ED, et al. Human extrafollicular CD4+ Th cells help memory B cells produce Igs. J Immunol. 2018 Sep 1;201(5):1359–1372. doi: 10.4049/jimmunol.1701217. Epub 2018 Jul 20. PMID: 30030323; PMCID: PMC6112860.

13. Xie MM, Dent AL. Unexpected Help: Follicular regulatory T cells in the germinal center. Front Immunol. 2018 Jul 2;9:1536. doi: 10.3389/fimmu.2018.01536. PMID: 30013575; PMCID: PMC6036241.

14. Gensous N, Charrier M, Duluc D, Contin-Bordes C, Truchetet ME, Lazaro E, et al. T follicular helper cells in autoimmune disorders. Front Immunol. 2018 Jul 17;9:1637. doi: 10.3389/fimmu.2018.01637. PMID: 30065726; PMCID: PMC6056609.

15. Quinn JL, Axtell RC. Emerging role of follicular T helper cells in multiple sclerosis and experimental autoimmune encephalomyelitis. Int J Mol Sci. 2018 Oct 19;19(10):3233. doi: 10.3390/ijms19103233. PMID: 30347676; PMCID: PMC6214126.

16. Cao G, Chi S, Wang X, Sun J, Zhang Y. CD4+CXCR5+PD-1+ T follicular helper cells play a pivotal role in the development of rheumatoid arthritis. Med Sci Monit. 2019 Apr 25;25:3032–3040. doi:10.12659/MSM.914868. PMID: 31019190; PMCID: PMC6498883.

17. Chen X, Li W, Zhang Y, Song X, Xu L, Xu Z, et al. Distribution of peripheral memory T follicular helper cells in patients with schistosomiasis Japonica. PLoS Negl Trop Dis. 2015 Aug 18;9(8):e0004015.doi: 10.1371/journal.pntd.0004015. PMID: 26284362; PMCID: PMC4540279.

18. Zhang Y, Wang Y, Jiang Y, Pan W, Liu H, Yin J, et al. T follicular helper cells in patients with acute schistosomiasis. Parasit Vectors. 2016 Jun 6;9(1):321. doi:10.1186/s13071-016-1602-6. PMID: 27266984; PMCID: PMC4895967.

19. Chen X, Yang X, Li Y, Zhu J, Zhou S, Xu Z, et al. Follicular helper T cells promote liver pathology in mice during Schistosoma japonicum infection. PLoS Pathog. 2014 May 1;10(5):e1004097. doi: 10.1371/journal.ppat.1004097. PMID: 24788758; PMCID: PMC4006917.

20. Wikenheiser DJ, Stumhofer JS. ICOS co-stimulation: friend or foe? Front Immunol. 2016 Aug 10;7:304. doi: 10.3389/fimmu.2016.00304. PMID: 27559335; PMCID: PMC4979228.

21. Le KS, Amé-Thomas P, Tarte K, Gondois-Rey F, Granjeaud S, Orlanducci F, et al. CXCR5 and ICOS expression identifies a CD8 T-cell subset with TFH features in Hodgkin lymphomas. Blood Adv. 2018 Aug 14;2(15):1889–1900. doi: 10.1182/bloodadvances.2018017244. PMID: 30087107; PMCID: PMC6093730.

22. Liu J, Zhou Y, Yu Q, Zhao Z, Wang H, Luo X, et al. Higher Frequency of CD4+CXCR5+ICOS+PD1+ T follicular helper cells in patients with infectious mononucleosis. Medicine (Baltimore). 2015 Nov;94(45):e2061. doi: 10.1097/MD.0000000000002061. PMID: 26559315; PMCID: PMC4912309.

23. Jogdand GM, Sengupta S, Bhattacharya G, Singh SK, Barik PK, Devadas S. Inducible costimulator expressing T cells promote parasitic growth during blood stage *plasmodium berghei* ANKA infection. Front Immunol. 2018 May 28;9:1041. doi: 10.3389/fimmu.2018.01041. PMID: 29892278; PMCID: PMC5985291.

24. Wikenheiser DJ, Ghosh D, Kennedy B, Stumhofer JS. The costimulatory molecule ICOS regulates host Th1 and follicular Th Cell differentiation in response to plasmodium chabaudi chabaudi AS infection. J Immunol. 2016 Jan 15;196(2):778–91. doi: 10.4049/jimmunol.1403206. Epub 2015 Dec 14. PMID: 26667167; PMCID: PMC4705592.

25. Tesciuba AG, Shilling RA, Agarwal MD, Bandukwala HS, Clay BS, Moore TV, et al. ICOS costimulation expands Th2 immunity by augmenting migration of lymphocytes to draining lymph nodes. J Immunol. 2008 Jul 15;181(2):1019–24. doi: 10.4049/jimmunol.181.2.1019. PMID: 18606653; PMCID: PMC2560985.

26. Shilling RA, Clay BS, Tesciuba AG, Berry EL, Lu T, Moore TV, et al. CD28 and ICOS play complementary non-overlapping roles in the development of Th2 immunity in vivo. Cell Immunol. 2009;259(2):177–84. doi:10.1016/j.cellimm.2009.06.013. Epub 2009 Jul 10. PMID: 19646680; PMCID: PMC2748173.

27. Ahamed J, Laurence J. Role of platelet-derived transforming growth factor-β1 and reactive oxygen species in radiation-induced organ fibrosis. Antioxid Redox Signal. 2017 Nov 1;27(13):977–988. doi: 10.1089/ars.2017.7064. Epub 2017 Jul 5. PMID: 28562065; PMCID: PMC5649128.

28. Jee MH, Hong KY, Park JH, Lee JS, Kim HS, Lee SH, et al. New mechanism of hepatic fibrogenesis: hepatitis C virus infection induces transforming growth factor β1 production through glucose-regulated protein 94. J Virol. 2015 Dec 30;90(6):3044–55. doi: 10.1128/JVI.02976-15. PMID: 26719248; PMCID: PMC4810663.

